# PNLDC1 catalysis and postnatal germline function are required for piRNA trimming, LINE1 silencing, and spermatogenesis in mice

**DOI:** 10.1101/2023.12.26.573375

**Authors:** Chao Wei, Xiaoyuan Yan, Jeffrey M. Mann, Ruirong Geng, Huirong Xie, Elena Y. Demireva, Liangliang Sun, Deqiang Ding, Chen Chen

**Affiliations:** Department of Animal Science, Michigan State University, East Lansing, Michigan 48824, USA; Shanghai Key Laboratory of Maternal and Fetal Medicine, Clinical and Translational Research Center, Shanghai First Maternity and Infant Hospital, Frontier Science Center for Stem Cell Research, School of Life Sciences and Technology, Tongji University, Shanghai 200092, China; Transgenic and Genome Editing Facility, Institute for Quantitative Health Science & Engineering, Michigan State University, East Lansing, Michigan 48824, USA; Department of Chemistry, Michigan State University, East Lansing, Michigan 48824, USA; Reproductive and Developmental Sciences Program, Michigan State University, East Lansing, Michigan 48824, USA; Department of Obstetrics, Gynecology and Reproductive Biology, Michigan State University, Grand Rapids, Michigan 49503, USA

## Abstract

PIWI-interacting RNAs (piRNAs) play critical and conserved roles in transposon silencing and gene regulation in the animal germline. Two distinct piRNA populations are present during mouse spermatogenesis: pre-pachytene piRNAs in fetal/neonatal testes and pachytene piRNAs in adult testes. PNLDC1 is required for both pre-pachytene piRNA and pachytene piRNA 3’ end maturation in multiple species. However, whether PNLDC1 is the bona fide piRNA trimmer and the physiological role of 3’ trimming of two distinct piRNA populations in spermatogenesis remain unclear. Here, by inactivating *Pnldc1* exonuclease activity in vitro and in mice, we reveal that PNLDC1 trimmer activity is required for both pre-pachytene piRNA and pachytene piRNA 3’ end trimming and male fertility. Furthermore, conditional inactivation of *Pnldc1* in postnatal germ cells causes LINE1 transposon de-repression and spermatogenic arrest in mice. This indicates that pachytene piRNA trimming, but not pre-pachytene piRNA trimming, is essential for mouse germ cell development and transposon silencing. Our findings highlight the potential of inhibiting germline piRNA trimmer activity as a potential means for male contraception.

## INTRODUCTION

PIWI-interacting RNAs (piRNAs) are recognized for their critical and conserved roles in silencing transposons and regulating genes within the germline, which are crucial for maintaining fertility across animal species (Czech et al., 2018; Iwasaki et al., 2015; Ozata et al., 2019; Siomi et al., 2011; Wang et al., 2023). These small RNAs, bound by PIWI proteins, function as guides in RNA-induced silencing complexes to regulate gametogenesis (Aravin et al., 2006; Girard et al., 2006; Grivna et al., 2006; Lau et al., 2006; Malone and Hannon, 2009). In mammals, disruptions in the piRNA pathway in germ cells are directly linked to spermatogenic impairments and consequent male infertility (Aravin et al., 2007; Carmell et al., 2007; Castañeda et al., 2011; Deng and Lin, 2002; Kuramochi-Miyagawa et al., 2004; Newkirk et al., 2017).

During mammalian spermatogenesis, two distinct piRNA populations are expressed at different germ cell developmental stages. In mice, the initial wave, known as fetal piRNAs or pre-pachytene piRNAs, is characterized by a high enrichment of transposon sequences. These piRNAs are associated with MILI and MIWI2 and play a critical role in transposon silencing, thereby safeguarding the integrity of the germline genome (Aravin et al., 2008; Aravin et al., 2007; Kuramochi-Miyagawa et al., 2008; Siomi et al., 2011). Conversely, at the onset of meiosis, specifically during the pachytene stage, a unique and abundant group of piRNAs exclusive to mammals emerges, known as pachytene piRNAs. They are produced from distinct intergenic piRNA clusters and differ significantly from their fetal counterparts in that they are poor in transposon sequences (Li et al., 2013; Özata et al., 2020; Sun et al., 2022). Pachytene piRNAs, in association with MILI and MIWI, are instrumental in regulating the differentiation and maturation of meiotic and post-meiotic germ cells. This regulation is pivotal for the successful completion of spermatogenesis and the production of viable spermatozoa (Castañeda et al., 2014; Choi et al., 2021; Dai et al., 2019; Ding et al., 2018; Goh et al., 2015; Gou et al., 2014; Vourekas et al., 2012; Wu et al., 2020; Zheng and Wang, 2012).

The production of both pre-pachytene piRNAs and pachytene piRNAs necessitates a conserved piRNA biogenesis machinery, which efficiently processes long single-stranded piRNA precursors through sequential steps, resulting in the formation of mature piRNAs associated with PIWI proteins (Iwasaki et al., 2015; Mohn et al., 2015; Ozata et al., 2019; Watanabe et al., 2011). In mammalian male germ cells, piRNA precursor processing predominantly occurs within the intermitochondrial cement (IMC), also known as nuage or germ granules (Aravin et al., 2009; Lehtiniemi and Kotaja, 2018; Wang et al., 2020; Yabuta et al., 2011). This process involves a sequence of endonucleolytic and exonucleolytic cleavages, resulting in the formation of mature PIWI-loaded piRNAs, which are typically 24-32 nucleotides in length. During this process, after pre-piRNAs are loaded onto PIWI proteins, a crucial trimming step occurs, where the 3’ end of the pre-piRNAs is shortened through 3’-5’ exonuclease activity (Ding et al., 2017; Izumi et al., 2016; Kawaoka et al., 2011; Mann et al., 2023; Nishimura et al., 2018; Tang et al., 2016; Zhang et al., 2017). The final stage of piRNA biogenesis involves the addition of a 2’-O-methyl group to the 3’ end of trimmed piRNAs by methyltransferase HENMT1. This modification confers stability to the piRNAs, thereby preparing them for the protective and regulatory functions within the germline (Gainetdinov et al., 2021; Kirino and Mourelatos, 2007; Lim et al., 2015).

In both mice and silkworms, PARN-like domain containing 1 (PNLDC1) is required for the 3’ trimming of pre-piRNAs (Ding et al., 2017; Izumi et al., 2016; Nishimura et al., 2018; Zhang et al., 2017). As a member of the PARN family of RNases, PNLDC1 possesses a conserved CAF1 exonuclease domain. PNLDC1 interacts with TDRKH, and the absence of either PNLDC1 or TDRKH results in piRNA 3’ end extension in both species (Ding et al., 2019; Ding et al., 2017; Honda et al., 2013; Izumi et al., 2016; Saxe et al., 2013). In mice, PNLDC1 is essential for male fertility, with its loss leading to spermatogenic arrest at the elongated spermatid stage, culminating in azoospermia (Ding et al., 2017; Nishimura et al., 2018; Zhang et al., 2017). Recent research has reported various *PNLDC1* variants in human azoospermic patients, suggesting a monogenic cause of human male infertility (Fang et al., 2023; Nagirnaja et al., 2021; Sha et al., 2022; Wang et al., 2022). These mutations lead to an increase in piRNA length due to defective trimming activity of PNLDC1. In the context of piRNA maturation, both pre-pachytene piRNAs in prospermatogonia and pachytene piRNAs in postnatal germ cells undergo crucial 3’ end trimming. The absence of PNLDC1 leads to elongation in both piRNA populations (Ding et al., 2017; Nishimura et al., 2018). However, it is unclear how these individual piRNA population defects contribute to spermatogenic arrest observed in *Pnldc1* knockout mice. Notably, the loss of PNLDC1 results in the de-repression of LINE1 in adult male germ cells but not in prospermatogonia (Ding et al., 2017). This discrepancy raises further questions whether the elongated pre-pachytene piRNAs retain their functional capacity in LINE1 transposon silencing, thus necessitating deeper investigation into the roles of PNLDC1 in the piRNA pathway.

In this study, we generated *Pnldc1* catalytic mutant mice to elucidate the physiological role of PNLDC1 exonuclease activity *in vivo*. Our findings demonstrate that PNLDC1 catalysis is required for piRNA trimming and spermatogenesis, proving PNLDC1 is indeed the piRNA trimmer in mice. Additionally, conditional deletion of *Pnldc1* in postnatal germ cells in mice disrupts LINE1 transposon silencing and spermatogenesis, highlighting the importance of pachytene piRNA maturation in promoting spermatogenesis.

## RESULTS

### E30A mutation ablates the exonuclease activity of mouse PNLDC1 *in vitro*

PARN-like domain containing 1 (PNLDC1) contains a conserved CAF1 exonuclease domain with a key DEDD (Asp-Glu-Asp-Asp) motif for catalytic activity (Fig. 1A). Mutation of the DEDD motif causes PNLDC1 exonuclease inactivation in silkworm (Izumi et al., 2016). To test whether mouse PNLDC1 has exonuclease activity *in vitro* and whether it is catalyzed by the CAF1 nuclease domain, we mutated the 30^th^ Glu to Ala in the DEDD motif of mouse PNLDC1 (PNLDC1 E30A) (Fig. 1A) (Izumi et al., 2016). We first examined the expression and localization of ectopically expressed wildtype PNLDC1 and PNLDC1 E30A in HeLa cells. When singly expressed, Flag-tagged PNLDC1 (Flag-PNLDC1) and Flag-PNLDC1 E30A exhibited similar expression level and subcellular localization pattern (Fig. 1B). When co-expressed with GFP-tagged TDRKH (TDRKH-GFP), mitochondrial-anchored TDRKH recruited both PNLDC1 and PNLDC1 E30A to the mitochondria (Fig. 1B). This indicates that PNLDC1 E30A mutation does not affect mutant protein expression, stability, and its recruitment to mitochondria by TDRKH to participate in piRNA trimming. To test the trimmer activity of PNLDC1, we next established a robust *in vitro* PNLDC1 trimming assay by combining the 293T cell-based heterologous expression system with an *in vitro* RNA trimming assay. In this system, Flag-PNLDC1 and TDRKH-GFP were co-expressed in 293T cells and PNLDC1-TDRKH complexes were immunoprecipitated by Flag-antibody conjugated beads to incubate with synthesized RNA substrates to detect RNA trimming. We showed that PNLDC1 alone was unable to trim RNA substrates but displayed strong trimming activity in the presence of TDRKH (Fig. 1C). This is consistent with previous reported PNLDC1 activity in the silkworm (Izumi et al., 2016). Strikingly, PNLDC1 E30A mutation completely disrupts the PNLDC1 trimming activity without affecting its interaction with TDRKH (Fig. 1C). We confirmed this finding using two independent RNA oligonucleotides as substrates (Fig. 1C). Together, these data demonstrate that mouse PNLDC1 has trimmer activity and E30A mutation abolishes its exonuclease activity *in vitro*.

**Fig 1.**
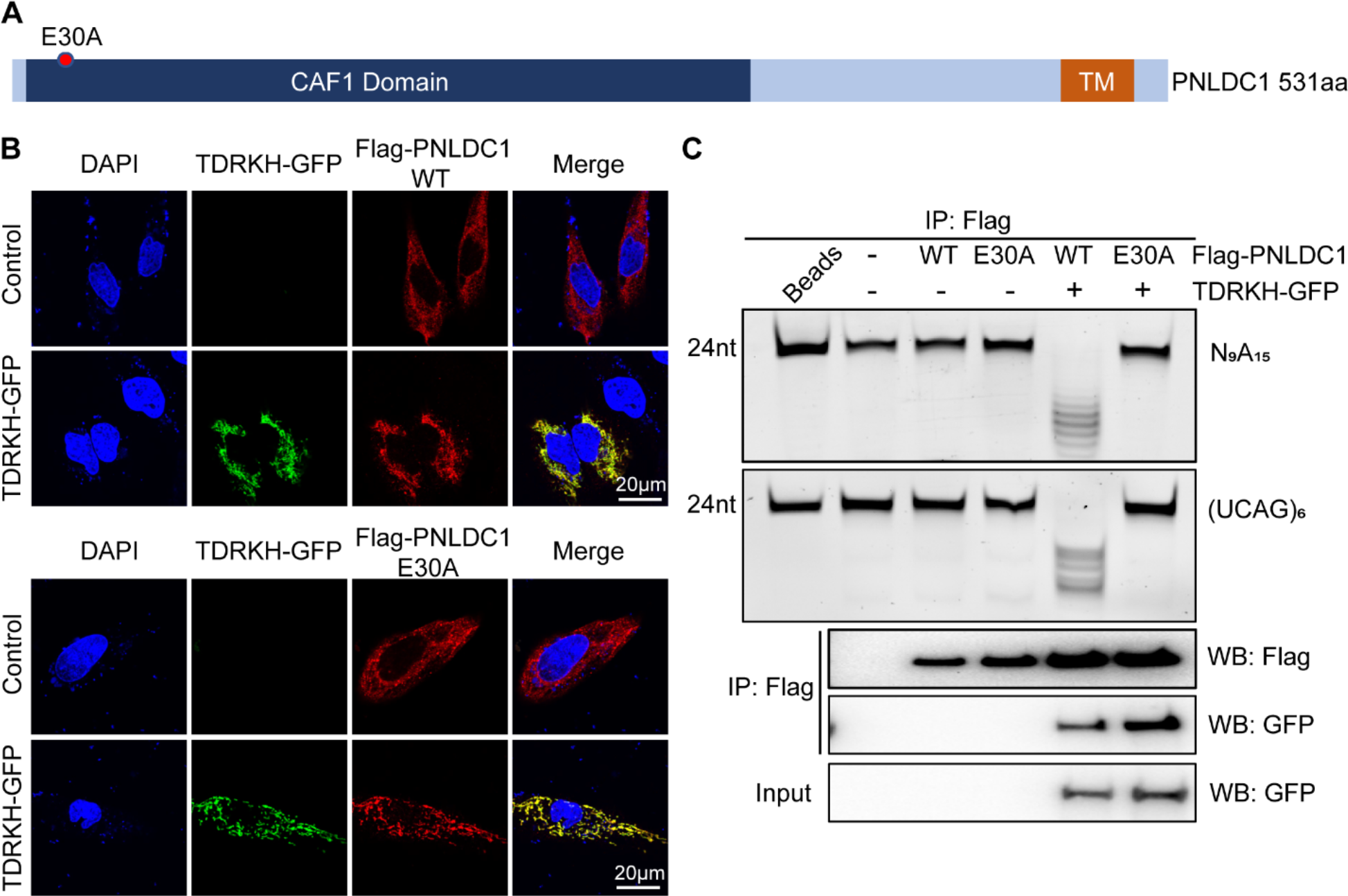
E30A mutation disrupts the exonuclease activity of PNLDC1 *in vitro*. **(A)** shows a schematic diagram of PNLDC1 protein domains and E30A mutation. **(B)** TDRKH recruits PNLDC1 and PNLDC1 E30A to mitochondria. HeLa cells were transfected with Flag-tagged PNLDC1/PNLDC1 E30A alone or together with TDRKH-GFP plasmids. After 48 h, the cells were fixed and stained with anti-Flag antibody. DNA was stained with DAPI. Scale bar, 20 μm. **(C)** 293T cells were transfected with plasmids expressing wild-type or catalytically inactive (E30A) PNLDC1, and the cell lysates were immunoprecipitated and analyzed by Western blotting (bottom). *In vitro* trimming assay was performed to detect the exonuclease activity of wild-type or catalytically inactive (E30A) PNLDC1. Results shown in (B) and (C) are representative of 3 biological replicates.

### PNLDC1 exonuclease inactivation causes LINE1 upregulation and male infertility in mice

To investigate whether PNLDC1 trimmer activity is essential for piRNA trimming and spermatogenesis *in vivo*, we generated a *Pnldc1* E30A mutant allele (*Pnldc1^E30A^*) in mice using CRISPR/Cas9 genome editing (Fig. S1A). The mutation in *Pnldc1^E30A^* mice was confirmed by Sanger DNA sequencing (Fig. S1B). We further generated *Pnldc1^E30A/-^*mice by breeding the *Pnldc1^E30A^* allele into the *Pnldc1* null (*Pnldc1^-^*) background (Ding et al., 2017). *Pnldc1^E30A/-^* mice are viable and grow normally but exhibited mildly reduced testis mass compared to *Pnldc1^+/-^*control littermates (Fig. 2A and B). This reduction in testis weight is similar to that observed in *Pnldc1^-/-^*testes (Ding et al., 2017). Histological analysis of spermatogenesis revealed a spermatogenic arrest at the elongated spermatid stage in *Pnldc1^E30A/-^*testes, a spermatogenic failure that phenocopies *Pnldc1^-/-^* mice (Fig. 2C and D). As a result, in *Pnldc1^E30A/-^* epididymis, only sloughed spermatids and residual cytoplasm were observed without normal spermatozoa being present, leading to male infertility (Fig. 2C).

**Fig 2.**
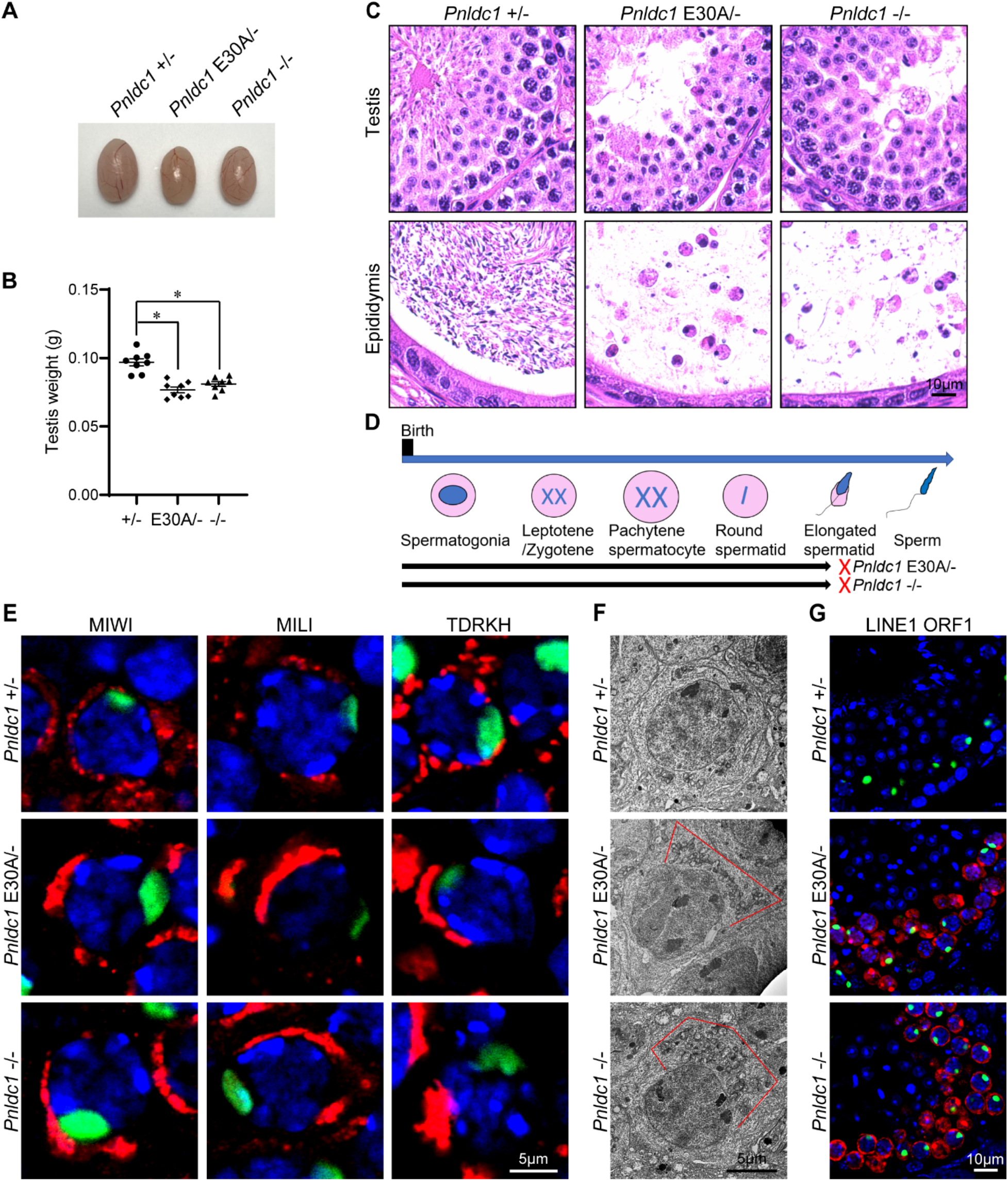
*Pnldc1* E30A mutation causes spermatocyte mitochondrial aggregation, LINE1 derepression, and spermatogenic arrest in mice. (A) Images of *Pnldc1^+/-^*, *Pnldc1^E30A/-^*, and *Pnldc1^-/-^* testes. **(B)** Testis weights of *Pnldc1^+/-^*, *Pnldc1^E30A/-^*, and *Pnldc1^-/-^* mice are shown. n = 8. Error bars represent SEM. The P-value was calculated by unpaired t-test. *, P < 0.01. **(C)** Hematoxylin and eosin-stained testis and epididymis sections from *Pnldc1^+/-^*, *Pnldc1^E30A/-^*, and *Pnldc1^-/-^* mice are shown. Scale bar, 10μm. **(D)** Spermatogenic arrest stage in *Pnldc1^E30A/-^* and *Pnldc1^-/-^* mice. **(E)** Immunostaining of MIWI, MILI, and TDRKH in *Pnldc1^+/-^*, *Pnldc1^E30A/-^*, and *Pnldc1^-/-^* spermatocytes. DNA is stained with DAPI. Scale bar, 5μm. **(F)** Transmission electron microscopy was performed on pachytene spermatocytes from *Pnldc1^+/-^*, *Pnldc1^E30A/-^*, and *Pnldc1^-/-^* testes. The mitochondria aggregation is indicated by red line. Scale bar, 5μm. **(G)** Immunostaining of LINE1 ORF1 in *Pnldc1^+/-^*, *Pnldc1^E30A/-^*, and *Pnldc1^-/-^* spermatocytes. DNA is stained with DAPI. Scale bar, 10μm. Results shown in (D) and (F)-(H) are representative of 3 biological replicates.

In adult testes, piRNA precursors are processed to load onto MIWI and MILI in intermitochondrial cements (IMCs) among mitochondrial clusters in pachytene spermatocytes. The piRNA biogenesis machinery comprises multiple factors including the PNLDC1/TDRKH trimming complex. We examined the expression and localization of MIWI, MILI, and TDRKH in *Pnldc1^E30A/-^* testes. Western blotting showed similar expression levels of TDRKH and MILI in *Pnldc1^+/-^*, *Pnldc1^E30A/-^*, and *Pnldc1^-/-^* testes (Fig. S2A). However, MIWI protein level was decreased in *Pnldc1^E30A/-^* testes, representing a similar MIWI reduction observed in *Pnldc1^-/-^* testes (Fig. S2A). Immunofluorescence showed that MIWI, MILI, and TDRKH localized to polarized and aggregated large perinuclear granules in *Pnldc1^E30A/-^*pachytene spermatocytes, suggesting mitochondrial aggregation in these cells (Fig. 2E and S2B). Transmission electron microscopy confirmed our observation: mitochondria in *Pnldc1^E30A/-^* spermatocytes displayed clustered aggregation and polar distribution, a defect also observed in *Pnldc1^-/-^*spermatocytes (Fig. 2F). Since PNLDC1 is required for LINE1 transposon silencing (Ding et al., 2017; Zhang et al., 2017), we next tested whether the trimmer activity of PNLDC1 is crucial for this role. By immunostaining, transposon LINE1 ORF1 protein was significantly upregulated in *Pnldc1^E30A/-^*, indicating the trimmer activity of PNLDC1 is indispensable for LINE1 suppression (Fig. 2G). Taken together, we conclude that PNLDC1 trimmer activity is required for LINE1 transposon silencing, spermiogenesis, and male fertility in mice.

### PNLDC1 exonuclease activity is essential for pachytene piRNA trimming

We next investigated the effect of PNLDC1 exonuclease inactivation on piRNA biogenesis. We used RNA labeling to examine the abundance and size of piRNA populations from adult *Pnldc1^+/-^*, *Pnldc1^E30A/-^*, and *Pnldc1^-/-^* testes. Radiolabeling of total RNA showed a normal piRNA population of around 30 nt in length in *Pnldc1^+/-^* testes (Fig. 3A). However, an abnormally longer but low abundant small RNA population of 30-40 nt in length was observed *Pnldc1^E30A/-^*testes, consistent with the extended piRNA population in *Pnldc1^-/-^*testes (Fig. 3A). We further performed MIWI and MILI immunoprecipitations and isolated MIWI-bound piRNAs (MIWI-piRNAs) and MILI-bound piRNAs (MILI-piRNAs) from *Pnldc1^+/-^*, *Pnldc1^E30A/-^*, and *Pnldc1^-/-^* testes. Radiolabeling assay showed that MIWI-piRNAs and MILI-piRNAs from *Pnldc1^E30A/-^* were similar to those from *Pnldc1^-/-^*, and both were longer than corresponding piRNAs in *Pnldc1^+/-^* (Fig. 3B and C). We further sequenced small RNA libraries constructed from total piRNA, MIWI-piRNAs and MILI-piRNAs in these mice and observed the same trend: a significant decrease in total piRNA amount (Fig. 3D) and an upshift piRNA lengths in *Pnldc1^E30A/-^* and *Pnldc1^-/-^*compared to *Pnldc1^+/-^* controls (Fig. 3D-F). Importantly, at the piRNA population level, piRNA defects of *Pnldc1^E30A/-^* and *Pnldc1^-/-^*were indistinguishable.

**Fig 3.**
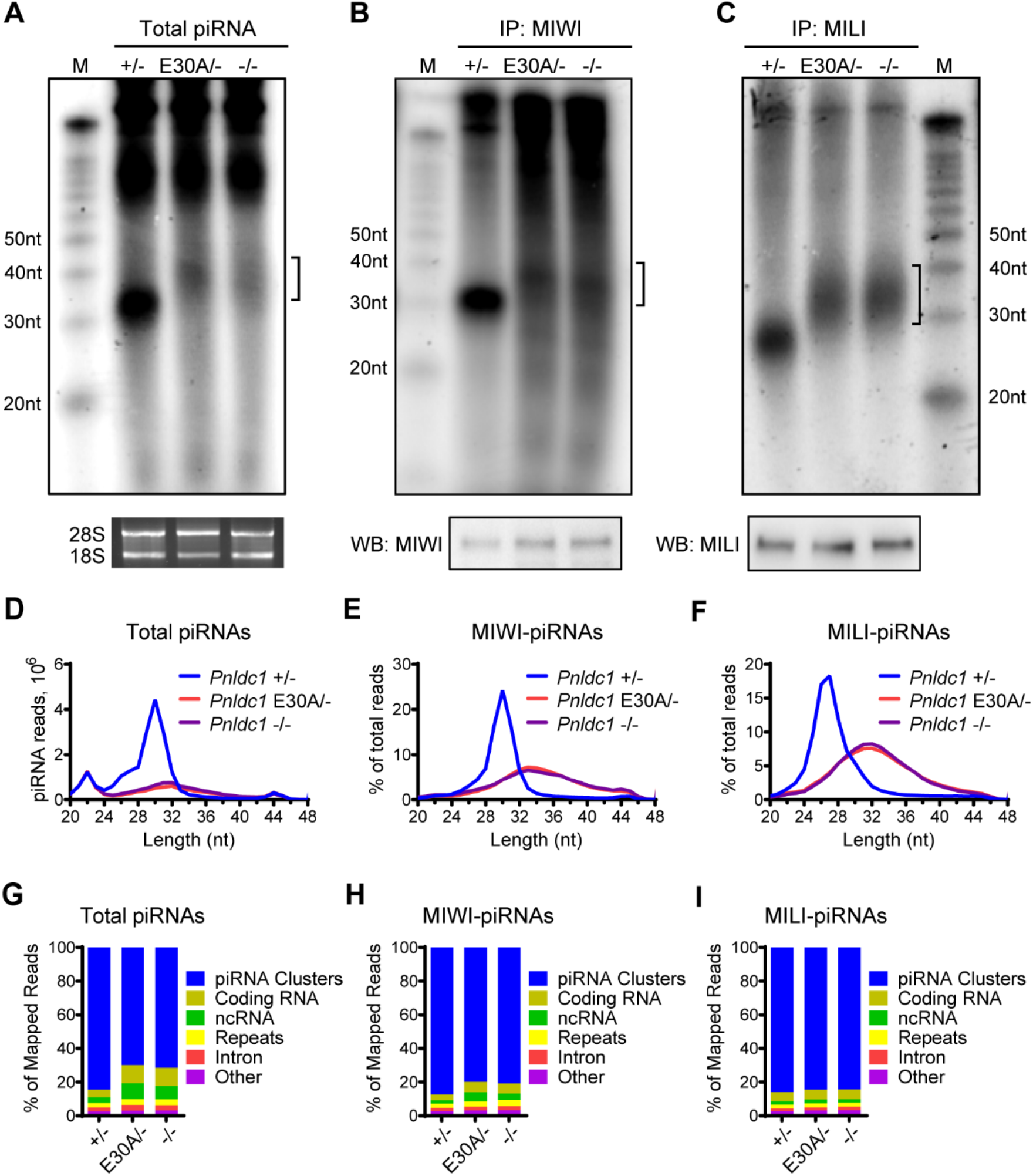
*Pnldc1* E30A mutation triggers defective pre-piRNA trimming. **(A)** Total RNAs extracted from *Pnldc1^+/-^*, *Pnldc1^E30A/-^*, and *Pnldc1^-/-^*testes were end-labeled with [^32^P]-ATP, separated by 15% TBE urea gel, and detected by autoradiography. Square bracket indicates extended piRNAs. The 18S and 28S ribosomal RNAs served as loading controls. **(B)** RNAs were isolated from immunoprecipitated MIWI RNPs and were end-labeled with [^32^P]-ATP, separated by 15% TBE urea gel, and detected by autoradiography. Western blotting was performed with anti-MIWI antibody to show immunoprecipitation efficiency. Square bracket indicates extended piRNAs. **(C)** RNAs were isolated from immunoprecipitated MILI RNPs and were end-labeled with [^32^P]-ATP, separated by 15% TBE urea gel, and detected by autoradiography. Western blotting was performed with anti-MILI antibody to show immunoprecipitation efficiency. Square bracket indicates extended piRNAs. **(D)** The length distribution of small RNAs from *Pnldc1^+/-^*, *Pnldc1^E30A/-^*, and *Pnldc1^-/-^*testicular total small RNA libraries. Data were normalized by microRNA reads (21–23 nt). **(E)** The length distribution of MIWI-bound piRNAs from *Pnldc1^+/-^*, *Pnldc1^E30A/-^*, and *Pnldc1^-/-^* MIWI-piRNA libraries. **(F)** The length distribution of MILI-bound piRNAs from *Pnldc1^+/-^*, *Pnldc1^E30A/-^*, and *Pnldc1^-/-^* MILI-piRNA libraries. **(G)** Genomic annotation of total piRNAs from *Pnldc1^+/-^*, *Pnldc1^E30A/-^*, and *Pnldc1^-/-^* testes. Sequence reads (24–48 nt) from total piRNA libraries were aligned to mouse genomic sequence sets in the following order: piRNA clusters, coding RNA, non-coding RNA, repeats, intron, and other. **(H)** Genomic annotation of MIWI-bound piRNAs from *Pnldc1^+/-^*, *Pnldc1^E30A/-^*, and *Pnldc1^-/-^* testes. Sequence reads (24–48 nt) from MIWI-piRNA libraries were aligned to mouse genomic sequence sets in the following order: piRNA clusters, coding RNA, non-coding RNA, repeats, intron, and other. **(I)** Genomic annotation of MILI-bound piRNAs from *Pnldc1^+/-^*, *Pnldc1^E30A/-^*, and *Pnldc1^-/-^* testes. Sequence reads (24–48 nt) from MILI-piRNA libraries were aligned to mouse genomic sequence sets in the following order: piRNA clusters, coding RNA, non-coding RNA, repeats, intron, and other. Results shown in (A)-(C) are representative of 3 biological replicates.

We further characterized the extended piRNAs in *Pnldc1^E30A/-^*testes by mapping the reads to the mouse genome. *Pnldc1^E30A/-^* piRNAs resembled *Pnldc1^-/-^*piRNAs in several features: A) mainly mapped to piRNA clusters (Fig. 3G, H and I); B) strong U bias at the first nucleotide position, which is the 5’ end signature of wild-type pachytene piRNAs (Fig. S3A); C) piRNA 3’ end extension (Fig. S3B). Collectively, these data demonstrate that the PNLDC1 exonuclease activity is required for pachytene pre-piRNA 3’ end trimming and therefore PNLDC1 is indeed the piRNA trimmer in mice.

### PNLDC1 exonuclease activity is required for pre-pachytene piRNA trimming

Next, we investigated the effect of PNLDC1 exonuclease inactivation on pre-pachytene piRNA biogenesis in prospermatogonia. By examining MILI and MIWI2 localization in neonatal (P0) *Pnldc1^+/-^*, *Pnldc1^E30A/-^*, and *Pnldc1^-/-^* testes, we found that MILI was expressed in a cytoplasmic granular pattern in prospermatogonia of all three mice (Fig. 4A). While MIWI2 was predominantly localized in the nuclei of *Pnldc1^+/-^* prospermatogonia, it only partially localized to nuclei with majority of signals in the cytoplasm of *Pnldc1^E30A/-^* and *Pnldc1^-/-^*prospermatogonia (Fig. 4B). The mis-localization of MIWI2 in *Pnldc1^E30A/-^* suggests defects in pre-pachytene piRNA biogenesis and function. Because pre-pachytene piRNAs are essential for LINE1 silencing, we examined transposon LINE1 expression in *Pnldc1^E30A/-^* prospermatogonia. Similar to *Pnldc1^+/-^*and *Pnldc1^-/-^*, no obvious LINE1 ORF1 was observed in *Pnldc1^E30A/-^*prospermatognia (Fig. 4C). This contrasts with the significant upregulation of LINE1 in *Tdrkh^-/-^* prospermatogonia and suggests that the remaining nuclear MIWI2 in *Pnldc1^E30A/-^* prospermatogonia is still functional for LINE1 silencing. We further explored whether PNLDC1 exonuclease inactivation affected pre-pachytene piRNA trimming by small RNA sequencing of MILI-piRNAs in neonatal (P0) testes. Similarly with results from adult testes, the peak of MILI-piRNAs shifted from 27 nt in *Pnldc1^+/-^* to 31-33 nt in *Pnldc1^E30A/-^* and *Pnldc1^-/-^* testes (Fig. 4D). Together, we conclude that PNLDC1 exonuclease activity is required for pre-pachytene piRNA trimming, but dispensable for LINE1 silencing in prospermatogonia.

**Fig 4.**
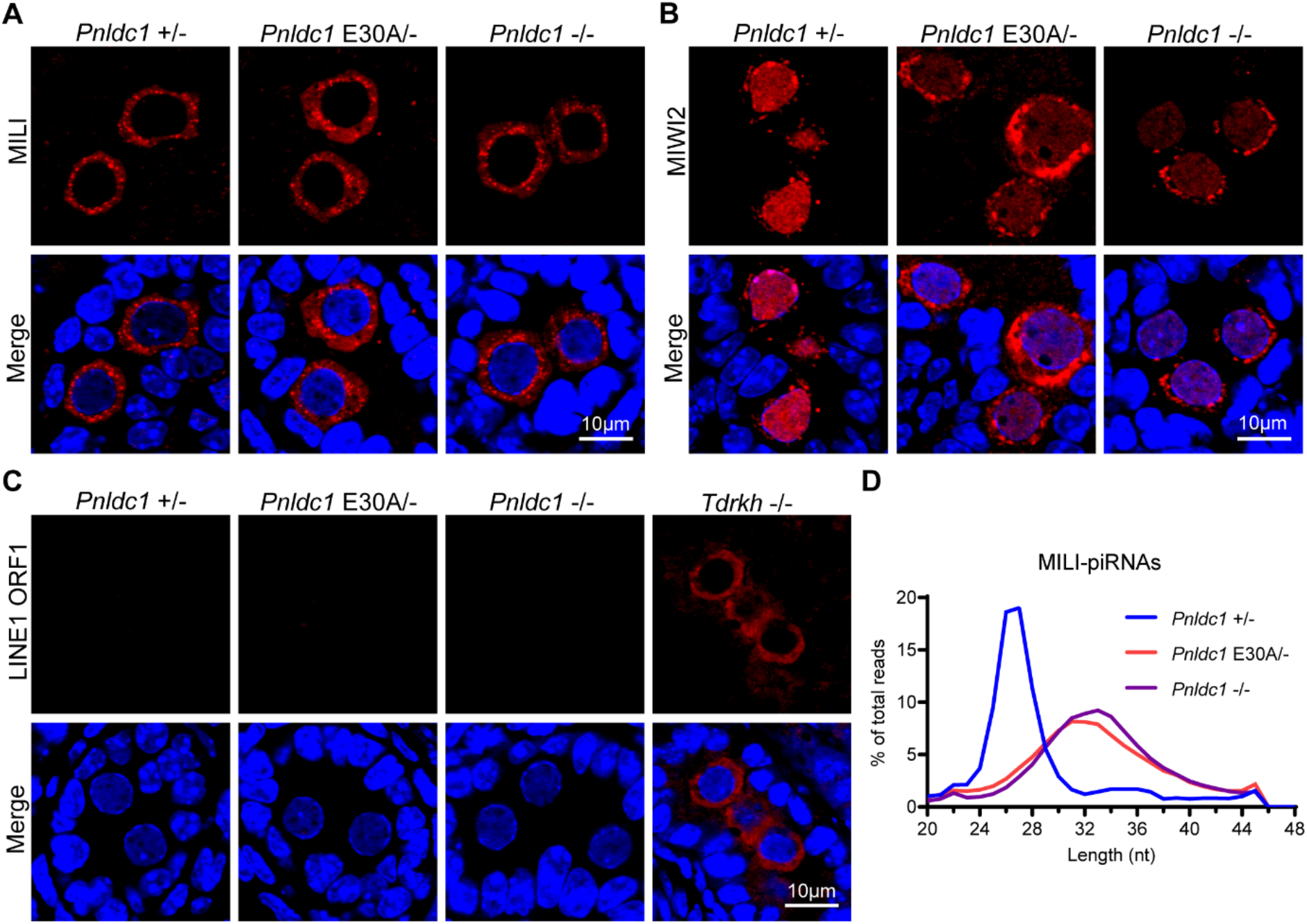
*Pnldc1* E30A mutation causes MIWI2 mislocalization but not LINE1 activation in neonatal prospermatogonia. **(A)** Immunostaining of MILI on neonatal (P0) testis sections from *Pnldc1^+/-^*, *Pnldc1^E30A/-^*, and *Pnldc1^-/-^* mice. Scale bar, 10μm. **(B)** Immunostaining of MIWI2 on neonatal (P0) testis sections from *Pnldc1^+/-^*, *Pnldc1^E30A/-^*, and *Pnldc1^-/-^* mice. Scale bar, 10μm. **(C)** Immunostaining of LINE1 ORF1 on neonatal (P0) testis sections from *Pnldc1^+/-^*, *Pnldc1^E30A/-^*, *Pnldc1^-/-^*, and *Tdrkh^-/-^* mice. Scale bar, 10μm. **(D)** The length distribution of MILI-bound piRNAs from neonatal (P0) *Pnldc1^+/-^*, *Pnldc1^E30A/-^*, and *Pnldc1^-/-^* MILI-piRNA libraries. Results shown in (A)-(C) are representative of 3 biological replicates.

### Postnatal germ cell-specific conditional deletion of *Pnldc1* in mice leads to LINE1 de-repression and spermatogenic arrest

It is unclear whether the spermatogenic arrest and transposon LINE1 de-repression in *Pnldc1^E30A/-^* and *Pnldc1^-/-^*testes is due to the defect of trimming in pre-pachytene piRNAs or pachytene piRNAs. To answer this, we generated a *Pnldc1* flox allele in mice using CRISPR-Cas9 genome editing (Fig. S4A). The exon 2 of *Pnldc1* was flanked by two loxP sites allowing its deletion by the Cre recombinase to ablate gene function. By combining *Pnldc1* flox with *Stra8*-Cre, we obtained *Stra8*-Cre^+^, *Pnldc1* flox/- conditional knockout (*Pnldc1* cKO) mice in which *Pnldc1* was deleted in male germ cells starting at postnatal day 3 (Fig. 5A). This deletion would not affect PNLDC1 expression during pre-pachytene piRNA biogenesis but only ablate PNLDC1 expression in postnatal germ cells. *Pnldc1* cKO mice are viable but exhibited reduced testis mass compared with control mice (Fig. 5B and C). Histological examination of *Pnldc1* cKO testes revealed that germ cells were primarily arrested at the elongated spermatid stage without normal spermatozoa in the epididymis, phenocopying *Pnldc1^-/-^* germ cell defect (Fig. 5D and E). These results indicate that postnatal function of PNLDC1 in germ cells is essential for spermatogenesis and male fertility.

**Fig 5.**
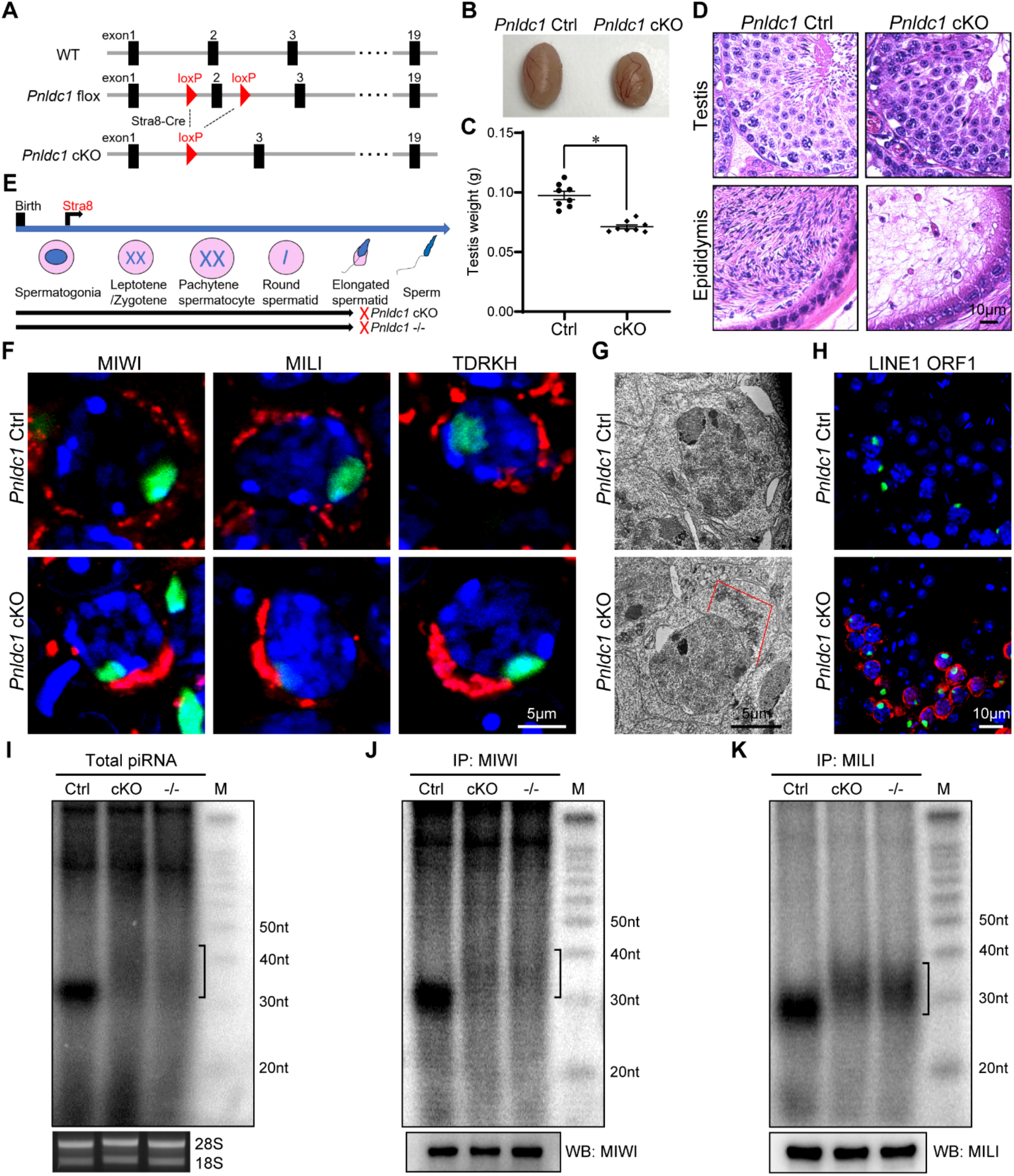
Conditional deletion of *Pnldc1* in postnatal germ cells causes defective pachytene piRNA trimming, LINE1 upregulation, and male infertility. **(A)** A schematic diagram showing the gene targeting strategy for the generation of *Pnldc1* ckO. Cre-mediated deletion removed the exon 2 of *Pnldc1*. **(B)** Images of control and *Pnldc1* cKO testes. **(C)** Testis weights of control and *Pnldc1* cKO mice are shown. n = 8. Error bars represent SEM. The P-value was calculated by unpaired t-test. *, P < 0.01. **(D)** Hematoxylin and eosin-stained testis and epididymis sections from control and *Pnldc1* cKO mice are shown. Scale bar, 10μm. **(E)** Spermatogenic arrest stage in *Pnldc1* cKO and *Pnldc1* KO mice. **(F)** Immunostaining of MIWI, MILI, and TDRKH in control and *Pnldc1* cKO spermatocytes. DNA is stained with DAPI. Scale bar, 5μm. **(G)** Transmission electron microscopy was performed on pachytene spermatocytes from control and *Pnldc1* cKO testes. The mitochondria aggregation is indicated by red line. Scale bar, 5μm. **(H)** Immunostaining of LINE1 ORF1 in control and *Pnldc1* cKO spermatocytes. DNA is stained with DAPI. Scale bar, 10μm. **(I)** Total RNAs extracted from control and *Pnldc1* cKO testes were end-labeled with [^32^P]-ATP, separated by 15% TBE urea gel, and detected by autoradiography. Square bracket indicates extended piRNAs. The 18S and 28S ribosomal RNAs served as loading controls. **(J)** RNAs were isolated from immunoprecipitated MIWI RNPs and were end-labeled with [^32^P]-ATP, separated by 15% TBE urea gel, and detected by autoradiography. Western blotting was performed with anti-MIWI antibody to show immunoprecipitation efficiency. Square bracket indicates extended piRNAs. **(K)** RNAs were isolated from immunoprecipitated MILI RNPs and were end-labeled with [^32^P]-ATP, separated by 15% TBE urea gel, and detected by autoradiography. Western blotting was performed with anti-MILI antibody to show immunoprecipitation efficiency. Square bracket indicates extended piRNAs. Results shown in (D) and (F)-(K) are representative of 3 biological replicates.

We further examined the expression and localization of MIWI, MILI, and TDRKH in *Pnldc1* cKO testes. Western blotting showed similar expression levels of TDRKH and MILI in control and *Pnldc1* cKO testes (Fig. S5A). However, MIWI protein level was decreased in *Pnldc1* cKO testes (Fig. S5A). Immunofluorescence showed that MIWI, MILI, and TDRKH formed polarized and aggregated large perinuclear granules in *Pnldc1* cKO pachytene spermatocytes (Fig. 5F and S5B). This coincides with observed mitochondrial aggregation in *Pnldc1* cKO spermatocytes by transmission electron microscopy (Fig. 5G). Interestingly, LINE1 ORF1 protein was significantly upregulated in *Pnldc1* cKO spermatocytes like *Pnldc1^-/-^* mice (Fig. 5H). This indicates that postnatally expressed PNLDC1 is essential for LINE1 silencing.

We next examined the effect of postnatal deletion of *Pnldc1* on piRNA biogenesis. Small RNA labeling revealed that total piRNA from *Pnldc1* cKO testes was longer than that from control testes (Fig. 5I). Labeling of MIWI-piRNAs and MILI-piRNAs confirmed the presence of longer piRNA populations in *Pnldc1* cKO testes, resembling the piRNA trimming defect in *Pnldc1^-/-^* testes (Fig. 5J and K). Together, these results suggest that proper pachytene piRNA trimming by PNLDC1 is required for LINE1 suppression and spermatogenesis.

## DISCUSSION

Pre-piRNA 3’ trimming catalyzed by the piRNA trimmer is a critical step during piRNA maturation in diverse species. Here, by inactivating the CAF1 nuclease domain of PNLDC1 *in vitro* and in mice we demonstrate that PNLDC1 is a bona fide mammalian pre-piRNA trimmer and this trimmer activity is essential for male fertility. Furthermore, postnatal deletion of *Pnldc1* in male germ cells disrupts spermatogenesis, highlighting the potential of blocking the piRNA trimmer activity in adult males for fertility regulation.

*Pnldc1^E30A/-^* and *Pnldc1^-/-^* mice exhibited identical piRNA and spermatogenic defects in all aspects we examined, which include: untrimmed pre-pachytene piRNAs and pachytene piRNAs at 3’ ends; normal LINE silencing in neonatal prospermatogonia but LINE1 de-repression in adult spermatocytes; adult spermiogenic arrest at the elongated spermatid stage. These results demonstrate that the physiological role of PNLDC1 is tightly coupled with its exonuclease activity in piRNA regulation and spermatogenesis.

Both PNLDC1 and TDRKH are required for piRNA trimming in mice (Ding et al., 2019; Ding et al., 2017; Nishimura et al., 2018; Saxe et al., 2013; Zhang et al., 2017). Notably, *Tdrkh^-/-^* mice show more severe defects in piRNA production and spermatogenesis than *Pnldc1^-/-^* mice, indicating that TDRKH has broader functions in piRNA regulation beyond piRNA 3’ end trimming (Saxe et al., 2013). Our study demonstrates that the PNLDC1 exonuclease domain is crucial for piRNA trimming *in vitro* and *in vivo*, proving PNLDC1 as a bona fide piRNA trimmer. However, *in vitro* trimming assay reveals that PNLDC1 cannot function alone, its trimmer activity depends on the presence of TDRKH. TDRKH is a mitochondrial transmembrane protein capable of recruiting PNLDC1 to mitochondria in cultured cells and we previously showed that it directly binds and recruits MIWI to the IMC for piRNA production (Chen et al., 2009; Chen et al., 2011; Ding et al., 2019; Wei et al., 2023). Therefore, we propose that TDRKH acts as a mitochondrial scaffold protein to simultaneously engage PNLDC1 and PIWI proteins to regulate piRNA trimming and PIWI function. This model can explain the phenotypic differences between *Tdrkh^-/-^* and *Pnldc1^-/-^* mice. Our findings clarify that PNLDC1 is the piRNA trimming enzyme while TDRKH is not. However, how PNLDC1 and TDRKH interact to form the piRNA trimming complex remains unknown and requires further biochemical and structural investigation.

We show that PNLDC1 catalysis is required for the 3’ trimming of both pre-pachytene piRNAs and pachytene piRNAs, two distinct piRNA populations from different germ cell developmental stages. However, defective pre-pachytene piRNA trimming and pachytene piRNA trimming have different effects on germ cell function. Without PNLDC1 catalytic activity, LINE1 silencing is normal in neonatal prospermatogonia but becomes defective in adult spermatocytes. This suggests that pre-pachytene piRNA trimming mediated by PNLDC1 is not required for LINE1 suppression in prospermatogonia. Consistent with this, we observed partial nuclear localization of MIWI2, suggesting that the residual untrimmed MIWI2-piRNAs are still functional in transcriptionally silencing LINE1. Given that gene knockouts of most piRNA biogenesis factors affecting pre-pachytene piRNAs causes LINE1 mis-regulation in prospermatogonia and results in early meiotic germ cell arrest (Carmell et al., 2007; Kuramochi-Miyagawa et al., 2004; Ma et al., 2009; Saxe et al., 2013; Shoji et al., 2009; Watanabe et al., 2011), the milder spermatogenic defect in *Pnldc1^E30A/-^* and *Pnldc1^-/-^* mice could result from defective pachytene piRNA maturation alone.

One important advance is the clarification of germ cell-specific function of PNLDC1 by conditional deletion of *Pnldc1* in postnatal germ cells. *Pnldc1^cKO^* mice exhibit identical LINE1 de-repression and spermatogenic arrest as *Pnldc1^-/-^* mice. This not only excludes its potential other functions beyond the germline but also reveals the requirement of postnatal pachytene piRNA trimming in male fertility. Recently multiple PNLDC1 variants with defective piRNA processing have been detected in human patients with azoospermia, establishing a monogenic cause of human infertility (Fang et al., 2023; Nagirnaja et al., 2021; Sha et al., 2022; Wang et al., 2022). Thus, based on our genetic dissection of mammalian PNLDC1 catalysis and germ cell specific function, we propose targeted inhibition of PNLDC1 enzymatic activity as a novel means for non-hormonal male contraception.

## Materials and Methods

### Ethics statement

All animal procedures were approved by the Institutional Animal Care and Use Committee of Michigan State University. All experiments with mice were conducted ethically according to the Guide for the Care and Use of Laboratory Animals and institutional guidelines.

### Mouse strains

*Pnldc1^E30A^* mutant mice were generated by CRISPR-Cas9 targeting of the mouse *Pnldc1* locus (ENSMUSG00000073460). Wild-type NLS-Cas9 protein, synthetic crRNA, tracrRNA, and single-stranded oligodeoxynucleotide (ssODN) donor template from Integrated DNA Technologies were used. Protospacer (N)20 and protospacer adjacent motif (PAM) sequence corresponding to the crRNA used was 5’-GGTCTGGATATAGAGTTCAC*-*AGG-3’. Donor ssODN in the reverse orientation had the following sequence: 5’- GGTAAGACAGTAATTAGTACCTGATCTGTTGGGGCCGAGACAAGTTTGAACGCAGACCTGTGAAAGCTATATCCAGACCTACGGGAGCACAAAACAGAC-3’. Synthetic crRNA and tracrRNA were incubated at 95 ℃ for 5 min and cooled down to form RNA heteroduplexes, which were then incubated with Cas9 protein for 5 min at 37 ℃ to preform ribonucleoprotein (RNP) complexes. RNPs were electroporated into C57BL/6 mouse zygotes using a Gene Editor electroporator (BEX CO., LTD, Tokyo, Japan) (Qin et al., 2015). Embryos were implanted into pseudo-pregnant recipients according to standard procedures. Gene editing of offspring was assessed using PCR, T7 Endonuclease I assay, and Sanger sequencing of the target region. Primers for *Pnldc1* E30A genotyping PCR are 5’-TGTACAGCTGCTTACCTCCT-3’ and 5’-AACAAAAACCAGCCCGCAG-3’. PCR products were digested with Hpy166II (R0616S, NEB).

For the generation of *Pnldc1* flox mice by CRISPR-Cas9 targeting, protospacer (N)20 and PAM sequences corresponding to crRNAs were 5′-GTACAGCTGCTTACCTCCTG-GGG-3′ for *Pnldc1* intron 1, and 5′-TGTCTGGTAGGGTCTAACTA-AGG-3′ for *Pnldc1* intron 2. Donor ssODN had the following sequence: 5’- CTGTTTTCCATGACTGTGTACAGCTGCTTACCTCGGATCCATAACTTCGTATAGCATACATTATACGAAGTTATCTGGGGCATTTGCAGTGTGAAAGGTGGTGCTTTGCTCTGTGCCTGAGGATTTTTGTCTGTTTTGTGCTCCCGTAGGTCTGGATATAGAGTTCACAGGTCTGCGTTCAAACTTGTCTCGGCCCCAACAGATCAGGTACTAATTACTGTCTTACCTCACCGGTGTCTCCTTAGATAACTTCGTATAGCATACATTATACGAAGTTATGAATTCTTAGACCCTACCAGACAAACATTGTGTCTGATCAG. To generate *Pnldc1* cKO mice, *Stra8*-Cre transgenic mice (017490, Jackson Laboratory) were bred with *Pnldc1* flox/flox mice using the strategy described in Fig. S4B. Primers for *Pnldc1* flox genotyping PCR are 5’-CCTGGGAACTGGTGTTTGGT-3’ and 5’-AGCCTATCAGCATTTGGCCA-3’. Primers for *Stra8*-Cre genotyping PCR are 5’-GTGCAAGCTGAACAACAGGA-3’ and 5’-AGGGACACAGCATTGGAGTC-3’. Primers for internal control in *Stra8*-Cre genotyping PCR are 5’-CTAGGCCACAGAATTGAAAGATCT-3’ and 5’-GTAGGTGGAAATTCTAGCATCATCC-3’ (Fig. S4C).

*Pnldc1^-/-^* and *Tdrkh^-/-^* mice were generated and genotyped as previously described (Ding et al., 2019; Ding et al., 2017).

### Plasmid construction

The full-length mouse *Pnldc1* and *Pnldc1^E30A^* mutant cDNAs were amplified by PCR and cloned into pcDNA3-Flag (Flag tag at N-terminus) expression vector. The full-length mouse *Tdrkh* cDNA was amplified by PCR and cloned into pEGFP-N1 (GFP tag at C-terminus) expression vector.

### Histology

Mouse testes and epididymides were collected and fixed in Bouin’s solution (HT10132, Sigma-Aldrich) overnight at 4 ℃ and embedded in paraffin. Histological sections were cut at 5 μm, dewaxed, rehydrated and stained with hematoxylin and eosin.

### Immunofluorescence

Mouse testes were fixed in 4% paraformaldehyde (PFA) in PBS overnight at 4 ℃ and embedded in paraffin. Testis sections were cut at 5 μm, dewaxed and rehydrated. Antigen retrieval was performed in Tris-EDTA buffer (pH 9.0) or sodium citrate buffer (pH 6.0). Testis sections were blocked in 5% normal goat serum (NGS) at room temperature (RT) for 30 min. Testis sections were then incubated with anti-MIWI (1:100; 2079, Cell Signaling Technology), anti-MILI (1:100; PM044, MBL), anti-TDRKH (1:100; 13528-1-AP, Proteintech), anti-LINE1 ORF1 (1:800), anti-MIWI2 (1:50; ab21869, Abcam) or FITC-conjugated mouse anti-γH2AX (1:500; 16-202A, Millipore) in 5% NGS at 37 ℃ for 2 h. After washing with PBS, sections were incubated with Alexa Fluor 555 goat anti-rabbit IgG (1:500; A21429, Thermo Fisher Scientific) at RT for 1 h and mounted using Vectashield mounting media with DAPI (H-1200, Vector Laboratories). Fluorescence was photographed using Fluoview FV1000 confocal microscope.

### Western blotting

Mouse testes were collected and homogenized in RIPA buffer (J63306-AP, Thermo Fisher Scientific) with protease inhibitor (A32965, Thermo Fisher Scientific). Protein lysates were separated by 4-20% polyacrylamide gels (4561096, Bio-Rad) and transferred to PVDF membranes. The membranes were blocked in 5% non-fat milk at RT for 30 min and subsequently incubated with primary antibodies in 5% non-fat milk overnight at 4 ℃. The primary antibodies used were anti-MIWI (1:1000; 2079, Cell Signaling Technology), anti-MILI (1:2000; PM044, MBL) anti-TDRKH (1:4000; 13528-1-AP, Proteintech), or HRP-conjugated mouse anti-β-actin (1:5000; A3854, Sigma-Aldrich). Membranes were washed with TBST and incubated with HRP-conjugated goat anti-rabbit IgG (1:5000; 1706515, Bio-Rad) at RT for 1 h followed by chemiluminescent detection with ECL Substrate (1705060, Bio-Rad).

### Transmission electron microscopy

Mouse testes were fixed in 2.5% glutaraldehyde in 0.1 M cacodylate buffer overnight at 4 ℃. After washing with 0.1 M cacodylate buffer, the testes were post-fixed in 1% osmium tetroxide in 0.1 M cacodylate buffer at RT for 2 h. The testes were then dehydrated in gradient series of acetone, infiltrated and embedded in Spurr’s resin. Ultrathin sections were cut at 70 nm and post-stained with uranyl acetate and lead citrate. Images were taken with JEOL 1400 Flash Transmission Electron Microscope (Japan Electron Optics Laboratory, Japan).

### Immunoprecipitation of piRNAs and proteins

Mouse testes were collected and homogenized using lysis buffer (20 mM HEPES pH 7.3, 150 mM NaCl, 2.5 mM MgCl2, 0.2 % NP-40, and 1 mM DTT) with protease inhibitor (A32965, Thermo Fisher Scientific) and RNase inhibitor (N2615, Promega). The lysates were pre-cleared using Protein A agarose beads (11134515001, Sigma-Aldrich) at 4 ℃ for 2 h. Anti-MIWI (2079, Cell Signaling Technology) or anti-MILI (PM044, MBL) antibody together with Protein A agarose beads were added to the lysates and incubated at 4 ℃ for 4 h. The beads were washed in lysis buffer 5 times. Immunoprecipitated RNAs were isolated from the beads using Trizol reagent (15596026, Thermo Fisher Scientific) for piRNA labeling or small RNA library construction. For protein detection, immunoprecipitated beads were boiled in protein loading buffer for 5 min. Western blotting of MIWI or MILI was performed as described above.

### Detection of piRNAs

Total RNA was extracted from mouse testes using Trizol reagent (15596026, Thermo Fisher Scientific). Total RNA or immunoprecipitated RNA (MIWI or MILI) was de-phosphorylated with Shrimp Alkaline Phosphatase (M0371, NEB) and end-labeled using T4 polynucleotide kinase (M0201, NEB) and [γ-32P] ATP (NEG002A250UC, PerkinElmer). The 32P-labeled RNA was separated by 15% Urea-PAGE gel. Radioactive signal was detected by exposing the gel on phosphorimager screen followed by scanning on the Typhoon scanner (GE Healthcare).

### Small RNA libraries and bioinformatics

Small RNA libraries from immunoprecipitated RNAs or total RNA were prepared using Small RNA Library Prep Kit (E7300, NEB) following the manufacturer’s instructions. Multiple libraries with different barcodes were pooled and sequenced with the Illumina HiSeq 4000 or NovaSeq 6000 platform (MSU Genomic Core Facility).

Sequenced reads were processed with fastx_clipper (http://hannonlab.cshl.edu/fastx_toolkit/index.html) to clip the sequencing adapter read-through. Clipped reads were filtered by length (24-48 nt) and aligned to the following sets of sequences: piRNA clusters, coding RNAs, non-coding RNAs, repeats, introns, and other. Alignments were performed with Bowtie (one base mismatch allowed). Repeats included classes of repeats as defined by RepeatMasker.

### Fixed cell immunofluorescence

HeLa cells were cultured in Chamber Slides (154534, Thermo Fisher Scientific) and transfected with indicated plasmids using Lipofectamine 2000 (11668019, Thermo Fisher Scientific). After 24 h, HeLa cells were fixed with 4% PFA for 15 min, incubated in 0.2% Triton X-100 in PBS for 20 min and blocked in 5% NGS at RT for 30 min. HeLa cells were then incubated with anti-FLAG antibody (1:100; F1804, Sigma-Aldrich) overnight at 4 ℃. After washing with PBS, HeLa cells were incubated with Alexa Fluor 555 goat anti-mouse IgG (1:200; A21422, Thermo Fisher Scientific) at RT for 1 h and mounted using Vectashield mounting media with DAPI (H-1200, Vector Laboratories). Fluorescence was photographed using Fluoview FV1000 confocal microscope.

### Trimming assay

HEK293T cells were transfected with indicated plasmids using Lipofectamine 2000 (11668019, Thermo Fisher Scientific). After 24 h, cells were lysed in trimming buffer (20 mM HEPES-KOH pH 7.0, 100 mM NaCl, 1.5 mM MgCl2, 0.1 mM EDTA, 0.1 mM DTT, protease inhibitor and RNase inhibitor) containing 1% Triton X-100 and immunoprecipitation was performed using anti-FLAG M2 agarose beads (A2220, Sigma-Aldrich). After immunoprecipitation, the beads were resuspended in 20 μl trimming buffer containing 0.5 uM 5’FAM labeled RNA oligo (N9A15: UGCCGCCCCAAAAAAAAAAAAAAA, or UCAG6:

UCAGUCAGUCAGUCAGUCAGUCAG) and incubated in the shaker at 37 ℃ for 1 h. Samples were then separated by 20% Urea-PAGE gel at 250 V for 1 h and the fluorescent signal was captured by the ChemiDoc System (Bio-Rad). For protein detection, immunoprecipitated beads were boiled in protein loading buffer for 5 min. Western blotting was performed as described above using anti-FLAG antibody (1:1000; F1804, Sigma-Aldrich) or anti-GFP antibody (1:10000; Ab290, Abcam) and secondary antibodies HRP-conjugated goat anti-mouse IgG (1:5000; 1706516, Bio-Rad) or goat anti-rabbit IgG (1:5000; 1706515, Bio-Rad).

## Statistical analysis

All data are mean ± SEM and all statistical analyses between groups were analyzed by unpaired t-test.

## Data availability

All sequencing data are deposited in the Sequence Read Archive of NCBI under the accession number PRJNA1055908 (adult *Pnldc1^+/-^*, *Pnldc1^E30A/-^*, and *Pnldc1^-/-^*; and neonatal *Pnldc1^E30A/-^*) and SRP095532 (neonatal *Pnldc1^+/-^* and *Pnldc1^-/-^*). All data are in the manuscript and/or supporting information files.

## Acknowledgment

We thank X. Cheng for critical reading of the manuscript, J. Hu and J. Ireland for sharing equipment. Deqiang Ding was supported by grants from National Natural Science Foundation of China (32070841 to DD), Natural Science Foundation of Shanghai (20ZR1460000 to DD), Chen Chen was supported by grants from National Institute of General Medical Sciences (R01GM132490 to CC), Eunice Kennedy Shriver National Institute of Child Health and Human Development (R01HD084494 to CC), and National Institute of Food and Agriculture (MICL02690 to CC).

## Author contributions

D.D. and C.C. designed research; C.W. performed library constructions, deep sequencing, histology and immunostaining with assistance from X.Y. and J.M.M.; R.G. and L.S. performed bioinformatics analysis and data analysis; H.X. and E.Y.D. generated *Pnldc1* E30A mutant mice and *Pnldc1* flox mice; C.W., D.D. and C.C. wrote the manuscript; C.C. and D.D. supervised the project.

## Competing financial interests

The authors have declared that no competing interests exist.

---

**Fig S1.**
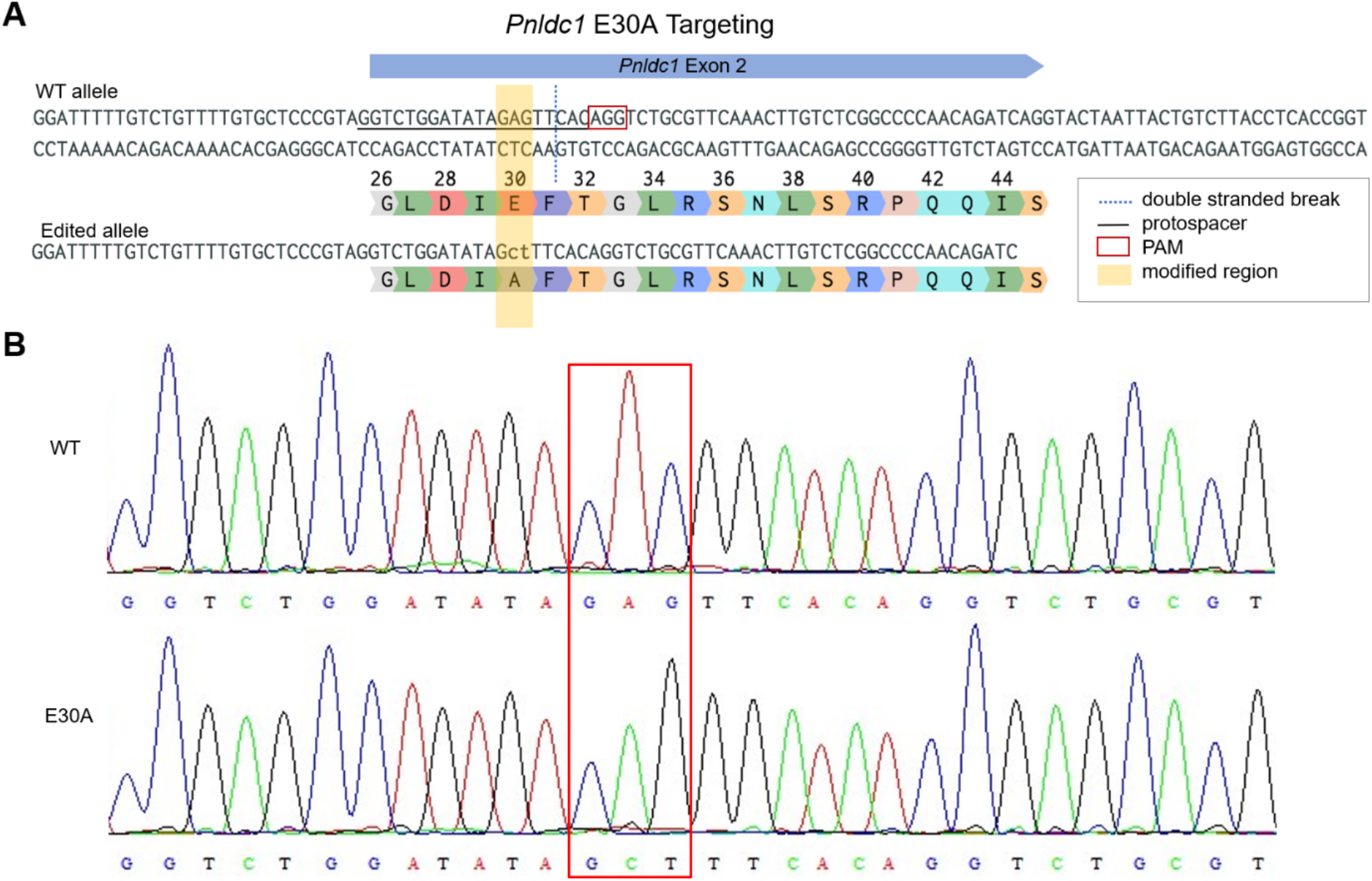
Targeting of the *Pndlc1* locus to generate E30A mutant mice. **(A)** The location of gRNA target protospacer and PAM, and the double stranded break following Cas9 cleavage are indicated on the WT allele. Modified codon E30A (GAG > Gct) is highlighted. The resulting edited allele sequence and translation are presented. **(B)** *Pnldc1* WT and E30A mutation are shown by Sanger DNA sequencing.

**Fig S2.**
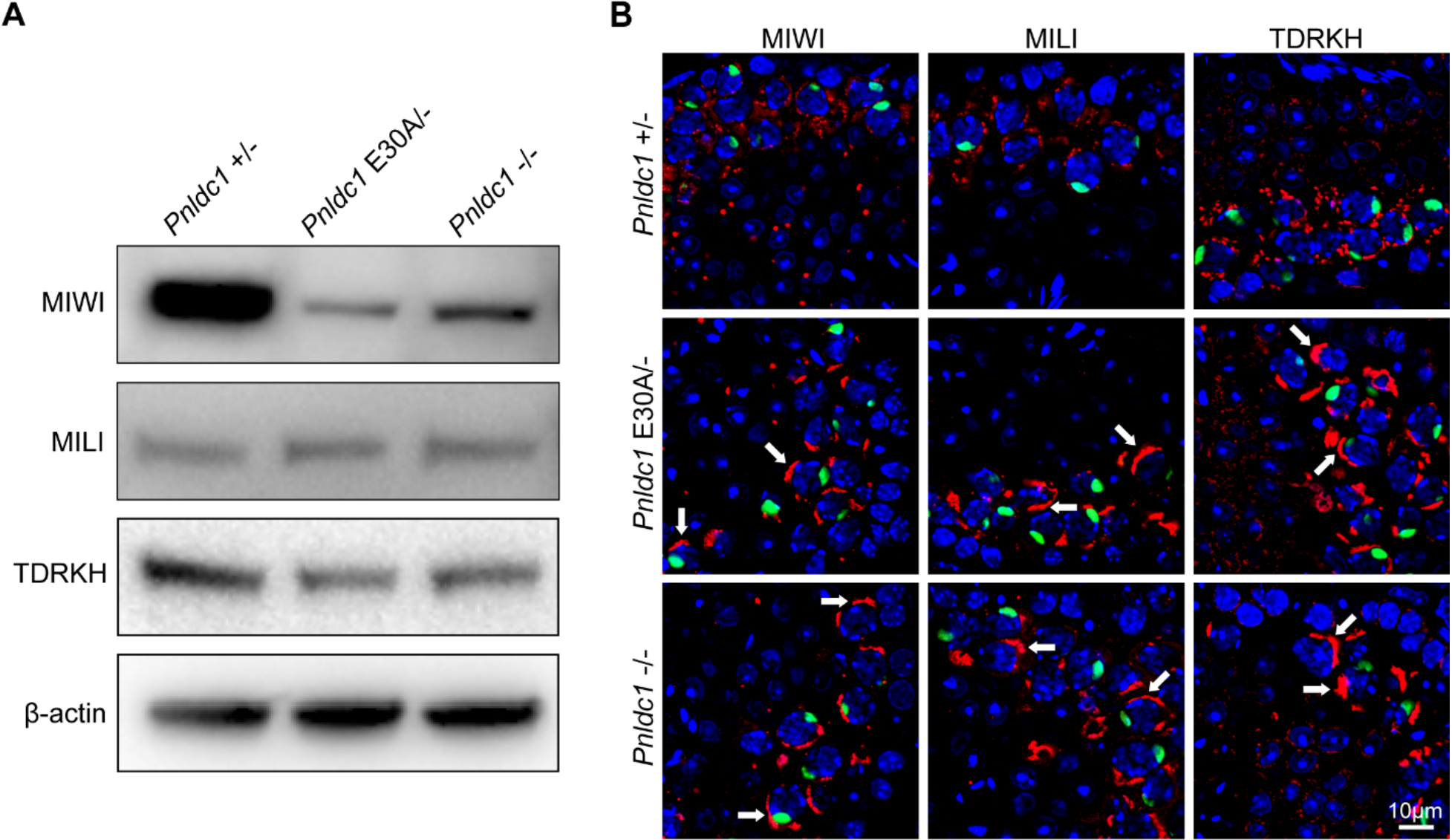
Expression and localization of MIW, MILI, and TDRKH in *Pnldc1^E30A/-^* testes. **(A)** Western blotting of MIWI, MILI, and TDRKH in *Pnldc1^+/-^*, *Pnldc1^E30A/-^*, and *Pnldc1^-/-^* testes. β-actin served as loading control. **(B)** Immunostaining of MIWI, MILI, and TDRKH in *Pnldc1^+/-^*, *Pnldc1^E30A/-^*, and *Pnldc1^-/-^* spermatocytes. DNA is stained with DAPI. Protein aggregation is indicated by arrows. Scale bar, 10μm. Results shown in (A) and (B) are representative of 3 biological replicates.

**Fig S3.**
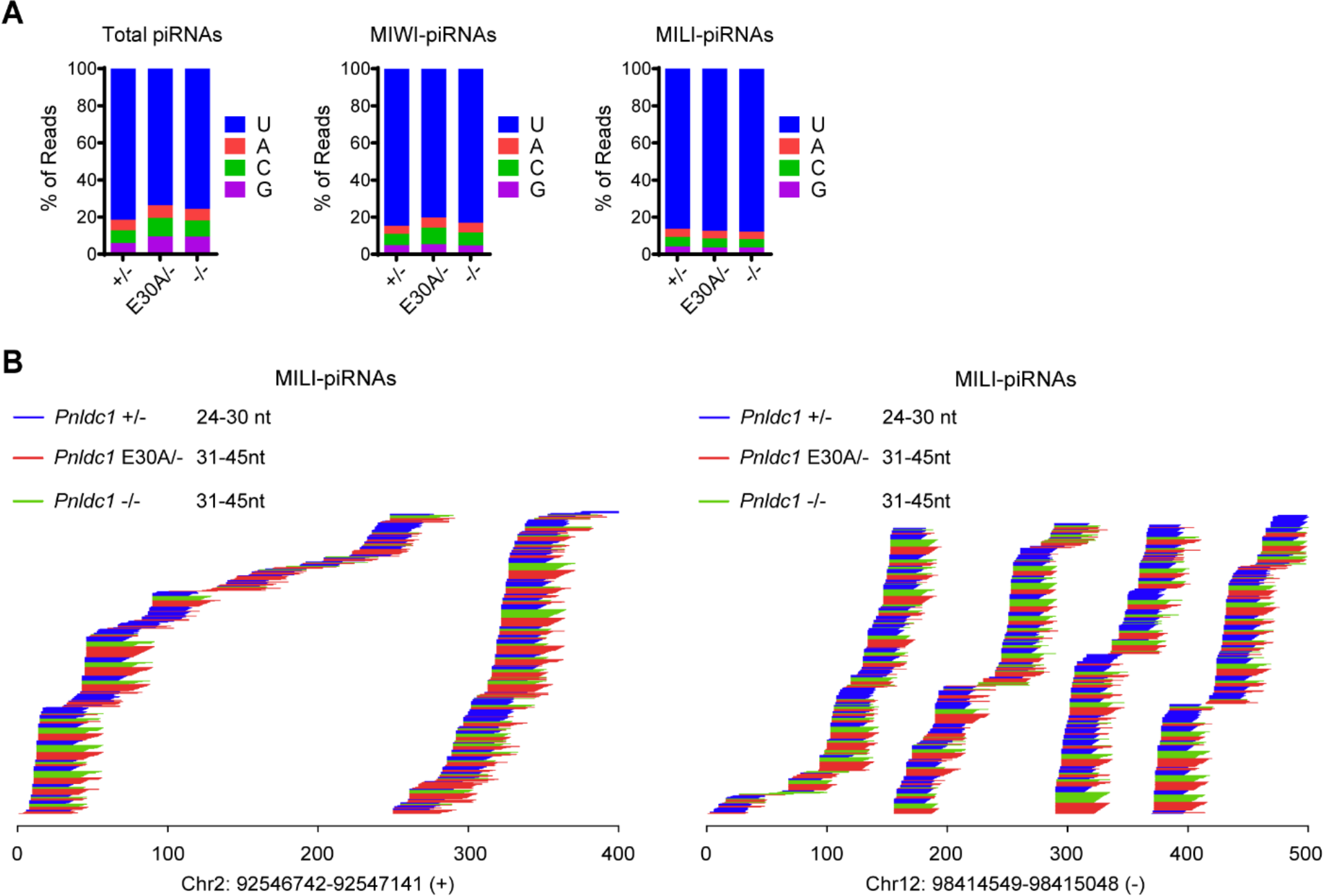
Extended piRNA 3’ ends in *Pnldc1^E30A/-^* testis. **(A)** Nucleotide distributions at the first position in total piRNA, MIWI-piRNAs, and MILI-piRNAs from *Pnldc1^+/-^*, *Pnldc1^E30A/-^*, and *Pnldc1^-/-^* testes. **(B)** Two examples of read alignments between MILI-piRNAs and piRNA clusters from *Pnldc1^+/-^*, *Pnldc1^E30A/-^*, and *Pnldc1^-/-^* testes. The genomic locations of the two piRNA clusters are shown at the bottom.

**Fig S4.**
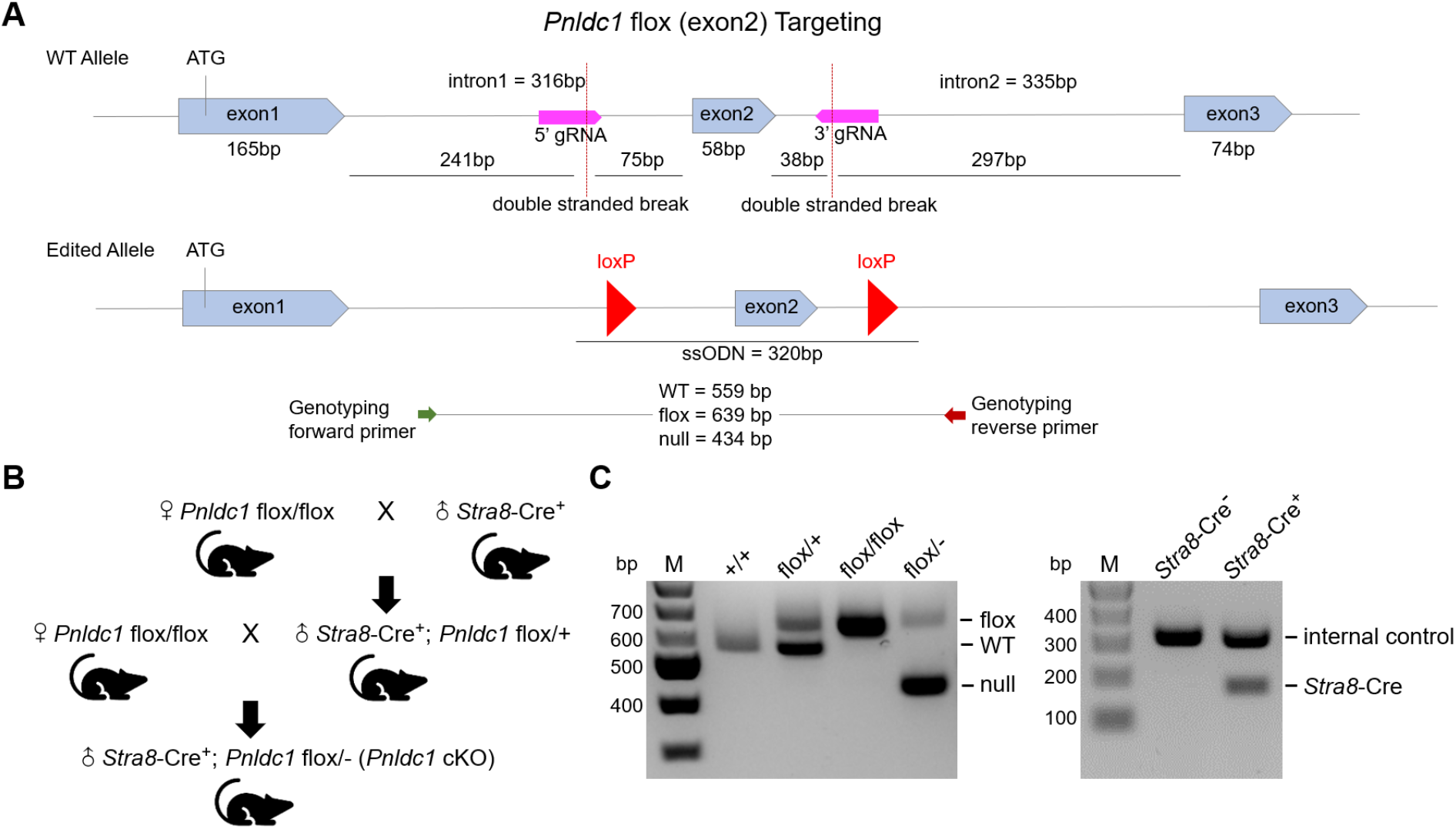
Targeting of the *Pndlc1* locus to generate *Pnldc1* flox mice. **(A)** The location of gRNAs and the double stranded break following Cas9 cleavage are indicated on the WT allele. The ssODN template containing two loxP sequences are indicated on the edited allele. **(B)** The breeding strategy to generate *Stra8*-Cre^+^; *Pnldc1* flox/- (*Pnldc1* cKO) mice. **(C)** Genotyping PCR of *Pnldc1* flox allele and *Stra8*-Cre allele. An internal control PCR was used to indicate the presence of genomic DNA.

**Fig S5.**
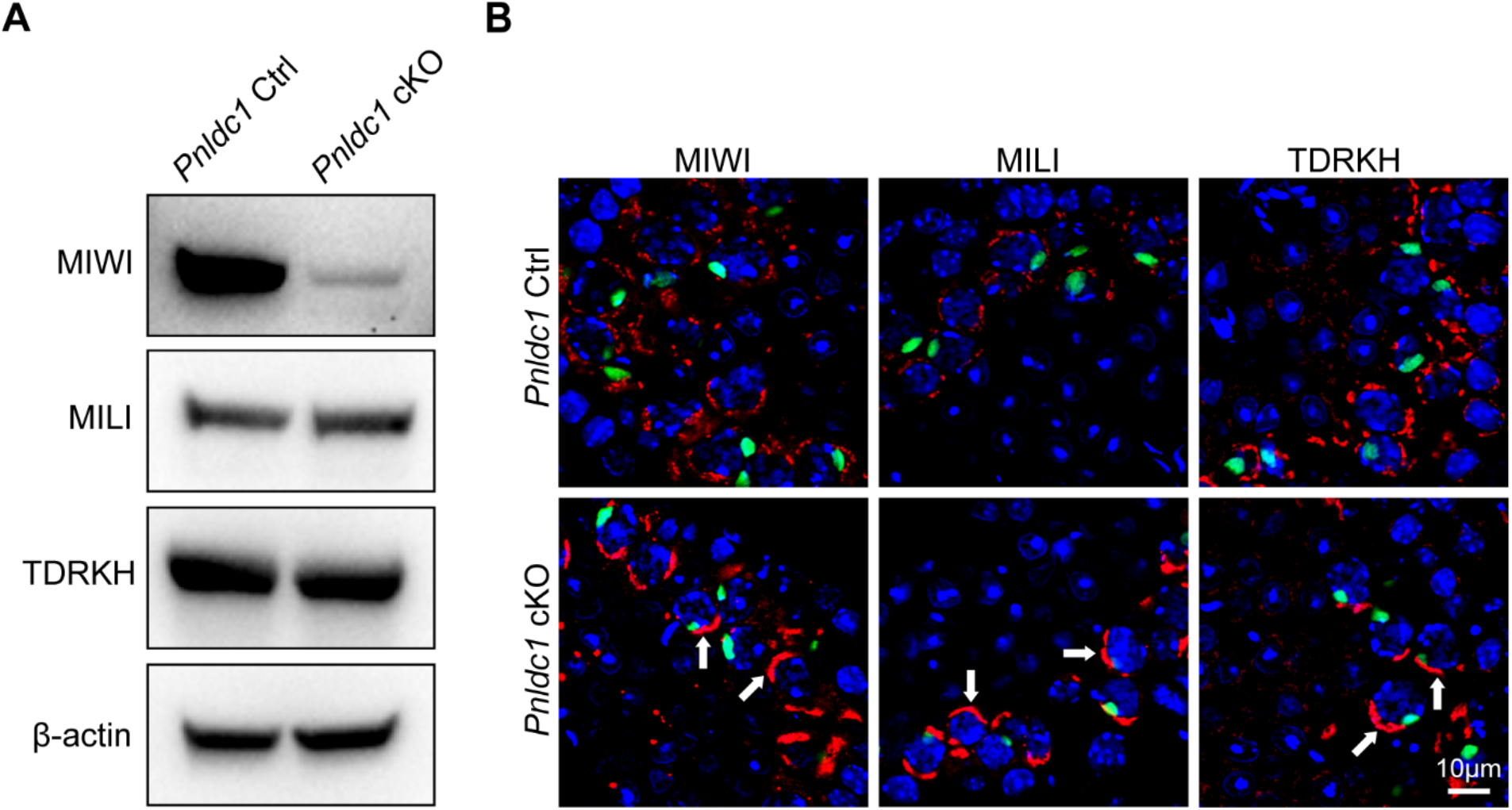
Expression and localization of MIW, MILI, and TDRKH in *Pnldc1* cKO spermatocytes. **(A)** Western blotting of MIWI, MILI, and TDRKH in control and *Pnldc1* cKO testes. β-actin served as loading control. **(B)** Immunostaining of MIWI, MILI, and TDRKH in control and *Pnldc1* cKO spermatocytes. DNA is stained with DAPI. Protein aggregation is indicated by arrows. Scale bar, 10μm. Results shown in (A) and (B) are representative of 3 biological replicates.

## Notes

### Competing Interest Statement

The authors have declared no competing interest.

